# Attending to what and where: Background connectivity integrates categorical and spatial attention

**DOI:** 10.1101/199190

**Authors:** Alexa Tompary, Naseem Al-Aidroos, Nicholas B. Turk-Browne

## Abstract

Top-down attention prioritizes the processing of goal-relevant information throughout visual cortex, based on where that information is found in space and what it looks like. Whereas attentional goals often have both spatial and featural components, most research on the neural basis of attention has examined these components separately. This may reflect the fact that attention is typically studied in individual visual areas that preferentially code for either spatial locations or particular features. Here we investigated how these attentional components are integrated by examining the attentional modulation of functional connectivity between visual areas with different selectivity. Specifically, we used fMRI to measure temporal correlations between spatially-selective regions of early visual cortex and category-selective regions in ventral temporal cortex while participants performed a task that benefitted from both spatial and categorical attention. We found that categorical attention modulated the connectivity of category-selective areas, but only with retinotopic areas that coded for the spatially attended location. The reverse was not true, however, with spatial attention modulating the connectivity of retinotopic areas with category-selective areas coding for both attended and unattended features. This pattern of results suggests that attentional modulation of connectivity is dominated by spatial selection, which in turn gates featural biases. Combined with exploratory analyses of frontoparietal areas that track these changes in connectivity among visual areas, this study begins to shed light on how different components of attention are integrated in support of more complex behavioral goals.

## Introduction

Top-down attention prioritizes the processing of goal-relevant information throughout the visual system. This enables the extraction of the most useful information from complex and noisy input. One neural mechanism that supports this prioritization is the amplified response of visual areas that code for goal-relevant information. Such amplified responses have been observed when attention is directed to particular spatial locations (Connor, Gallant, Preddie, & Essen, 1996; e.g., Moran & Desimone, 1985; Motter, 1993), and when it is directed to specific features, objects, and categories (e.g., Corbetta, Miezin, Dobmeyer, Shulman, & Petersen, 1990; O’Craven, Downing, & Kanwisher, 1999; Serences, Schwarzbach, Courtney, Golay, & Yantis, 2004; Furey et al., 2006). Another neural mechanism that supports prioritization is increased coupling between areas. Such increased coupling occurs among goal-relevant visual regions and with frontoparietal areas that control attention, again for both spatial (Bosman et al., 2012; Gregoriou, Gotts, Zhou, & Desimone, 2009; Griffis, Elkhetali, Burge, Chen, & Visscher, 2015; Saalmann, Pigarev, & Vidyasagar, 2007) and non-spatial attention (Al-Aidroos, Said, & Turk-Browne, 2012; Baldauf & Desimone, 2014; Zhou & Desimone, 2011).

Most of the studies above treat spatial and non-spatial attention as orthogonal in order to isolate their neural mechanisms. However, in real life, these two forms of attention are closely intertwined, such as when scanning the tables at a restaurant for a friend or when looking for a highway exit with a nearby gas station. This integration has been studied in the behavioral literature (e.g., Goldsmith & Yeari, 2003; Soto & Blanco, 2004), through response amplification in visual cortex (Müller & Kleinschmidt, 2003; e.g., Treue & Trujillo, 1999), and in terms of overlap in frontoparietal control areas (Giesbrecht, Woldorff, Song, & Mangun, 2003; Shomstein & Yantis, 2004, 2006). However, it is unclear how the integration of spatial and non-spatial attention is supported by coupling between areas.

In the current study, we examined changes in BOLD correlations between visual areas while attention is simultaneously allocated to both a location and a category. We presented a stream of faces and a stream of scenes, with one stream appearing to the left of fixation and the other to the right. Participants were instructed to detect repetitions in one stream, a task that benefitted from both spatial and categorical attention. We used a background connectivity approach to quantify attentional modulation of coupling. This approach is conceptually similar to resting-state connectivity (Fox & Raichle, 2007) except measured during tasks (Al-Aidroos et al., 2012; Duncan, Tompary, & Davachi, 2014; Griffis et al., 2015; Norman-Haignere, McCarthy, Chun, & Turk-Browne, 2012; Summerfield et al., 2006; Tompary, Duncan, & Davachi, 2015). We examined the background connectivity of V4 — a retinotopically organized occipital region that is modulated by attention (McAdams & Maunsell, 1999; Saenz, Buracas, & Boynton, 2002; Cohen & Tong, 2015) — with the fusiform face area (FFA) and the parahippocampal place area (PPA), ventral temporal regions sensitive to faces and scenes that are also modulated by attention (O’Craven et al., 1999; Serences et al., 2004).

This design differs from our previous study of categorical attention (Al-Aidroos et al., 2012), in which both face and scene streams were overlaid at one location. That study found stronger V4 correlations with FFA when faces were attended and with PPA when scenes were attended. By presenting the streams in different hemifields, we can exploit the contralateral organization of V4 to evaluate the role of spatial attention in this effect. On one hand, categorical attention might modulate coupling only between areas that code for spatially attended locations, similar to findings that spatial attention gates feature-based modulation of baseline activity (e.g., McMains, Fehd, Emmanouil, & Kastner, 2007). For example, when attending to faces on the right, FFA may show increased coupling only with left V4. On the other hand, coupling might be modulated by categorical attention even for areas that code for spatially unattended locations, similar to findings that feature-based modulation of responses occurs globally (e.g., Treue & Trujillo, 1999; Saenz et al., 2002; Andersen, Hillyard, & Müller, 2013). In the example above, FFA would also show increased coupling with right V4. Adjudicating between these hypotheses will help determine whether attentional modulation of coupling supports the integration of spatial and non-spatial attention.

## Materials and Methods

### Participants

Fifteen naïve right-handed members of the Princeton community (8 male, ages 18-35, mean = 23.6 years) participated for monetary compensation (20$/h). All participants had normal or corrected visual acuity and provided informed consent. The study protocol was approved by the Institutional Review Board for Human Participants at Princeton University.

### Overview of procedure

Each participant completed two experimental sessions while undergoing fMRI scanning and eye tracking. Both sessions were completed within 10 days of each other. In the first session, participants completed a repetition detection task in which they selectively attended to particular spatial locations and categories. We used this main session to investigate how different attentional states altered background connectivity in visual cortex. In the second session, participants completed a retinotopic mapping task followed by a face/scene localizer task. We used these tasks to identify regions of interest (ROIs) in occipital cortex (left and right V1-V4) and ventral temporal cortex (FFA and PPA), respectively

### First session

The first session contained six runs: two *rest* runs followed by four *attention* runs. During each rest run, participants were shown a white circular fixation point (radius = 0.2°) centered on a gray background for 441 s (Figure 1A). Participants were instructed to passively view the fixation point without performing any overt task. We used these rest runs to quantify baseline levels of connectivity.

**Figure 1.**
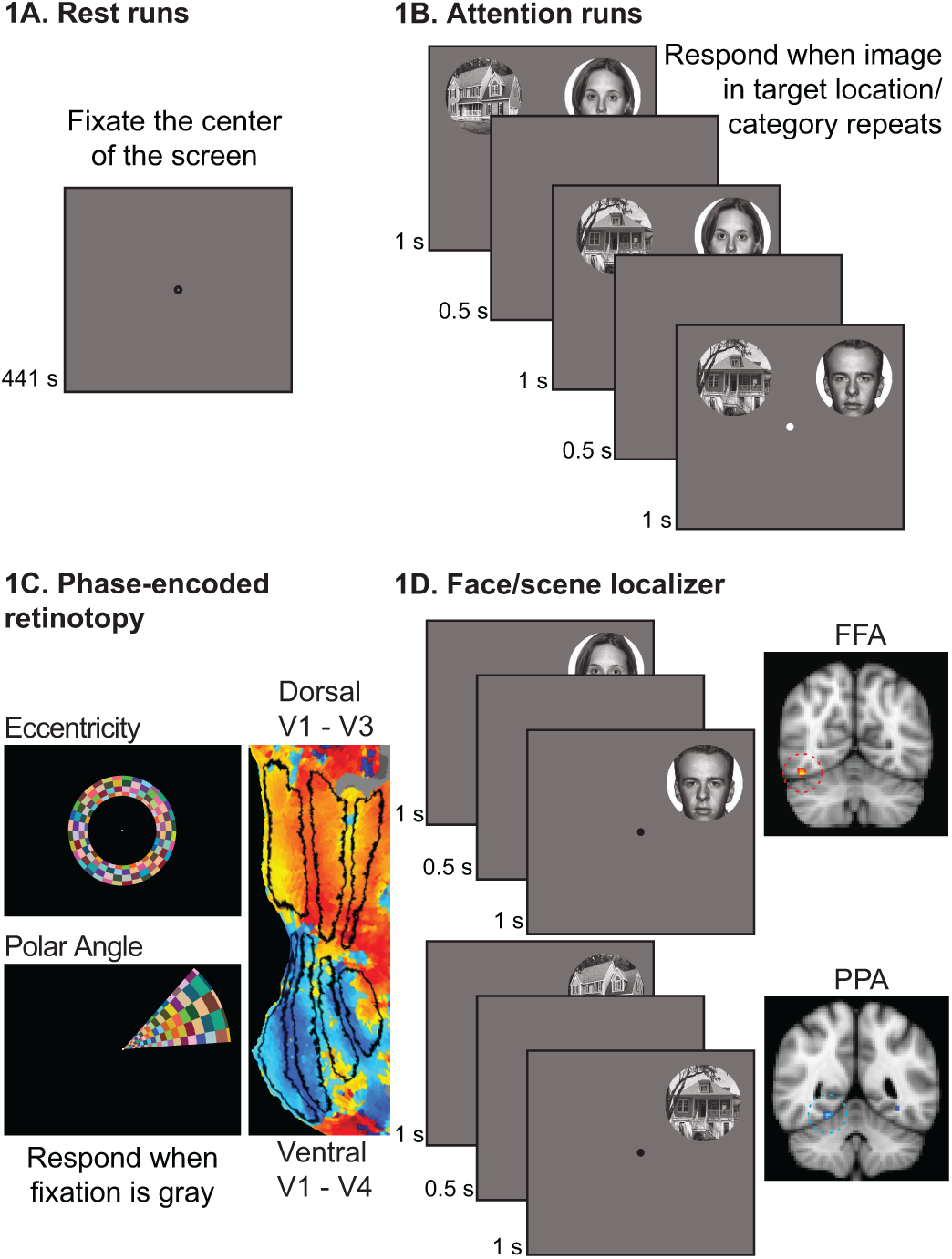
Experimental design. (A) In session 1, participants first completed two rest runs. They were instructed to fixate the dot in the center of the screen. (B) During attention runs, participants simultaneously viewed two streams of images and performed a repetition-detection task on one stream. Faces were presented in one stream and scenes in the other. The location and category of the attended stream was manipulated such that participants attended a unique combination of spatial locations and categories in each run: faces on the left, faces on the right, scenes on the left, and scenes on the right.(C) In session 2, participants completed six retinotopy scans: four polar scans with rotating wedges and two eccentricity scans with expanding/contracting rings. These scans were used to identify V4 and ventral/dorsal V1-V3. (D) Participants then completed two functional localizers for FFA and PPA. These runs were identical to the attention runs, except only one stream was presented at a time. The blocks within each scan alternated between face and scene stimuli.

The attention runs were used to assess background connectivity in different attentional states (Figure 1B). Attention runs started with 9 s of fixation followed by an on-off block design that alternated between twelve 18-s stimulation blocks and twelve 18-s fixation blocks (441 s total). Each stimulation block consisted of a sequence of twelve 1-s trials, separated by a 500-ms interstimulus interval. Each trial contained a face image and a scene image. Faces were drawn from a set of 24 photographs from the NimStim database (http://www.macbrain.org/resources.htm, neutral expressions). Scenes were drawn from a set of 24 photographs of single houses collected from the Internet and stock photograph CDs (Norman-Haignere et al., 2012). Images were converted to grayscale, cropped using a circular mask, and re-sized to a 6° radius. The two possible locations for the images were the left and right quadrants of the upper visual field (centered 8° above the horizontal meridian, and 8° to the left and right of the vertical meridian, respectively). These locations projected to the ventral stream in the visual processing hierarchy and to the contralateral right and left hemispheres, respectively.

Within each attention run, all images from a given category appeared in the same location (i.e., scenes on the left and faces on the right, or vice versa). We manipulated attentional state by instructing participants to perform a repetition detection task for one location and category, reporting whether the current task-relevant image was an immediate repeat of the last task-relevant image. Participants reported repetitions by pressing a button on a button box and otherwise withheld responses. To ensure that they attended to the entire stream of twelve images per block, task-relevant repetition targets occurred either once or twice unpredictably (split 50/50). In blocks with one repetition, the repetition randomly occurred at any point in the block. When repetitions occurred twice, they were separated by at least one non-repetition trial, and the second repetition happened within the final quarter of the block. To increase the need for selection, repetitions also occurred in the task-irrelevant location and category, but they were to be ignored. Otherwise, the images within a block were selected randomly from the respective category set without replacement.

Counterbalancing the location of face and scene images and the relevant category for the repetition detection task across attention runs resulted in a 2 (attention to face vs. scene category) x 2 (attention to left vs. right location) factorial design with four conditions: attention to faces in the left visual field, attention to faces in the right visual field, attention to scenes in the left visual field, and attention to scenes in the right visual field. Every participant completed one run of each type, in an order that was pseudo-counterbalanced across participants.

### Second session

The second session contained eight runs: six *retinotopy* runs followed by two *localizer* runs. The retinotopy runs were used to identify visual areas — left and right V1-V4 — containing topographic maps of the visual field. The stimuli and procedure were identical to the retinotopy task we have used previously (Al-Aidroos et al., 2012; see also, Arcaro, McMains, Singer, & Kastner, 2009). A dynamic and colorful circular checkerboard (4-Hz flicker, 14° radius) was shown at fixation to stimulate different visual receptive fields (Figure 1C). During *polar angle* runs, the checkerboard was masked so that only a 45° wedge was visible, and the mask rotated clockwise, or counterclockwise, at a rate of 9°/s (40 s period) creating the perception of a rotating wedge. These runs allowed us to identify the preferred polar angle of voxels in visual cortex. During *eccentricity* runs, the checkerboard was masked to create the perception of an expanding, or contracting, annulus. The expansion/contraction period was 40 s, and the thickness and rate of expansion of the annulus increased logarithmically with eccentricity to approximate the cortical magnification factor of early visual cortex (from 0.3-1.5°). These runs allowed us to identify the preferred eccentricity of stimulation for voxels in visual cortex. All participants completed one clockwise and one counterclockwise polar angle run (order counterbalanced), followed by one expanding and one contracting eccentricity run (order counterbalanced), and followed by a second clockwise and a second counter-clockwise polar angle run (same order as initial polar angle runs). To ensure that participants were actively processing the visual display, they were instructed to detect when a central fixation point (radius = 0.25°) dimmed from white to gray (every 2 to 5 s) using a button box.

The localizer runs were used to identify the FFA and PPA (Figure 1D). These runs were similar to the attention runs described above, but differed in two ways. First, we presented one image at a time, always in the left visual field for one run and in the right visual field for the other run (order counterbalanced). Second, the category of these images varied across blocks, alternating between faces and scenes within run (starting category randomized). In all other respects, the localizer runs were identical to the attention runs, including stimulus sizes, possible locations, photographs, trial timings, block timings, and task (i.e., repetition detection). For the first two participants, the localizer runs were completed at the end of the first session.

### Eyetracking

A critical component of the attention runs was that participants fixated a centrally presented point while covertly attending to peripheral spatial locations for the repetition-detection task. To be confident that stimuli fell in the anticipated retinotopic areas, we assessed fixation using a 60-Hz camera-based SMI iViewX MRI-LR eyetracker, mounted at the foot of the scanner bed. Calibration was performed at the beginning of each session and between runs when necessary. Drift correction was occasionally applied during fixation phases of the block (i.e., when only a fixation point was present). Data were recorded from whichever eye was tracked most accurately. The resulting gaze-direction timecourses were segmented into saccades, fixations, or noise (i.e., blinks or lost signal) using an algorithm similar to Nyström and Holmqvist (2010). Gaze direction could not be accurately recorded from 2/15 participants due to inconsistent pupil reflection or partially occluded pupils. We discarded eyetracking data from 7 runs total across all of the remaining participants because of poor or lost calibration. Within successfully recorded runs, 14.5% of data were discarded because of blinks and artifacts. We assessed participants’ ability to maintain fixation by calculating the average horizontal distance of gaze position from fixation and the percentage of samples more than 2° from fixation.

### Image acquisition

fMRI data were acquired with a 3T scanner (Siemens Skyra) using a 16-channel head coil. Functional images during rest, attention, and localizer runs were acquired with a gradient-echo echo-planar imaging sequence (repetition time [TR] = 1.5 s; echo time [TE] = 28 ms; flip angle [FA] = 64°; matrix = 64 × 64; resolution = 3 × 3 × 3.5 mm), with 27 interleaved axial slices aligned to the anterior-/posterior-commissure line. TRs were time-locked to the presentation of images (or the beginning of the run, in the case of rest runs). Functional images during retinotopy runs were acquired with a similar sequence but at higher spatial resolution (TR = 2.0 s; TE = 40 ms; FA = 71°; matrix = 128 × 128; resolution = 2 × 2 × 2.5 mm), with 25 interleaved slices aligned parallel to the calcarine sulcus.

For each session and functional sequence, we collected T1 fast low angle shot (FLASH) scans, aligned coplanar to the functional scan. Phase and magnitude field maps were acquired coplanar with the functional scans and with the same resolution, to correct B0-field inhomogeneities. Finally, a high-resolution magnetization-prepared rapid acquisition gradient-echo (MPRAGE) anatomical scan was acquired for surface reconstruction and registration.

### Image preprocessing

fMRI data were analyzed using FSL (Smith et al., 2004), FreeSurfer (Dale, Fischl, & Sereno, 1999; Fischl, Sereno, & Dale, 1999), and MATLAB (MathWorks). Preprocessing began by removing the skull from images to improve registration. Data from the first six volumes of functional runs were discarded for T1 equilibration. Remaining functional images were: motion corrected using MCFLIRT, corrected for slice-acquisition time, high-pass filtered with a 100-s period cutoff, debiased using the field maps in FUGUE, and spatially smoothed (retinotopy runs: 3-mm FWHM; all other runs: 5-mm FWHM). The processed images were registered to the high-resolution MPRAGE and Montreal Neurological Institute standard brain (MNI152). Retinotopy scans were first aligned to the FLASH to improve registration to the MPRAGE.

### Regions of interest

We identified FFA and PPA ROIs using the localizer runs. As in previous studies (Al-Aidroos et al., 2012; Norman-Haignere et al., 2012), we limited analyses to right FFA because the right hemisphere is dominant for face processing (Verosky & Turk-Browne, 2012; Yovel, Tambini, & Brandman, 2008). We limited analyses to right PPA to facilitate comparison with the right FFA, because its selectivity to spatial location is roughly equivalent (Schwarzlose, Swisher, Dang, & Kanwisher, 2008), although it likely exhibits less of a contralateral bias than FFA (Hemond, Kanwisher, & Op de Beeck, 2007). For each run, we fit a GLM with regressors for the face and scene blocks, modeled with 18-s boxcar regressors that were convolved using FSL’s double-gamma HRF. The temporal derivatives of these regressors were also included, as well as six regressors for different directions of head motion. We first contrasted parameter estimates for face vs. scene blocks within left- and right-visual field runs and then collapsed across visual field in a second-level GLM. We defined the FFA by choosing the voxel in right lateral fusiform cortex most selective for face stimuli (i.e., face > scene blocks) and defined the PPA by choosing the voxel in right collateral sulcus/parahippocampal cortex most selective for scene stimuli (i.e., scene > face blocks). In all analyses, BOLD signal from each ROI was extracted as a weighted average of the surrounding voxels, with weights determined by a Gaussian kernel with 5-mm FWHM centered on the peak voxel.

Based on the retinotopy runs, we drew ROIs in both hemispheres using FreeSurfer, including V4 and ventral and dorsal V1-V3. Retinotopy runs were pre-whitened, corrected for hemodynamic lag (3 s), and then phase decoded to determine the preferred polar angle and eccentricity of stimulated voxels in visual cortex. We further accounted for hemodynamic lag by averaging phase estimates from clockwise and counterclockwise runs, and from expansion and contraction runs. To facilitate ROI drawing, we plotted phase maps on 2-D surfaces segmented and flattened from the MPRAGE scans at the white matter/gray matter boundary. We identified boundaries between ROIs based on the relative locations of polar angle reversals and foveally stimulated regions (Arcaro et al., 2009; Wandell, Dumoulin, & Brewer, 2007).

### Evoked responses

We first assessed how attention modulated the amplitude of the BOLD responses in our ROIs. The preprocessed runs were fit with GLMs that captured the average evoked response in the attention blocks. We used a finite impulse response (FIR) model rather than a canonical hemodynamic response function because it avoided assumptions about the shape and timing of the response across voxels. Each model included 23 FIR regressors for the first 23 volumes of each block (including stimulation and fixation), with a delta function at the corresponding timepoint of all blocks and zeros elsewhere. The 24^th^ volume of each block (just before the next block) was left out to serve as a baseline. We also included six regressors for the motion correction parameters from preprocessing. The resulting parameter estimates were extracted from each ROI, transformed to percent signal change, and submitted to statistical tests.

### Background connectivity

To assess how attention modulated coupling, we examined correlations in the BOLD signal across ROIs for different attention runs. We used a background connectivity approach, which removed evoked responses and other noise sources that could induce spurious correlations. This approach has previously been used successfully to measure how shifts in categorical attention regulate the strength of interactions between regions of visual cortex (Al-Aidroos et al., 2012; Norman-Haignere et al., 2012). First, the preprocessed data were submitted to a ‘nuisance’ model. Specifically, we fit a GLM to each attention and rest run that included regressors for: the global mean BOLD signal, the six motion parameters from preprocessing, and the BOLD signal from four seeds in the ventricles and four seeds in white matter (left/right, anterior/posterior). The residuals of this model were submitted to a second ‘evoked’ GLM designed to remove evoked responses in a similar manner to above, except with an additional FIR regressor for the 24^th^ volume in each block (because we were interested in removing all evoked activity, rather than measuring responses relative to baseline). We measured background connectivity by extracting the residuals of this model from each ROI and relating them to each other with Pearson correlation. We applied Fisher’s r-to-z transformation to all coefficients before statistical tests. All reported correlations, including in the figures, are presented as the original *r* values to facilitate interpretation and visualization.

### Temporal MUD analysis

Finally, we conducted an exploratory analysis to identify potential control regions that could be responsible for modulating background connectivity in visual cortex. The measure of background connectivity described above provides a point estimate of the overall temporal relationship between brain areas. However, because none of the areas had perfect correlations, there was by definition variance in the extent to which individual timepoints supported this relationship. We reasoned that the activity of an additional area involved in controlling attention would fluctuate synchronously with this variance, with more activity (and greater modulation) during timepoints that strengthened the relationship among the other areas and less activity during timepoints that weakened it. This analysis involved two steps: (1) quantifying how much each timepoint contributed to background connectivity between visual ROIs, and (2) relating this variance to the activity timecourse of other voxels in the brain.

To quantify the contribution of each time point, we adapted the multivariate-univariate dependence (MUD) analysis technique, which was developed to measure how much each voxel contributed to a spatial correlation computed across multiple voxels (Aly & Turk-Browne, 2016b, 2016a). Switching from spatial to temporal correlation, we measured how much each timepoint contributed to the background connectivity previously computed across timepoints for each attention run. For example, timepoints when two visual areas were both active (or both inactive) would support a positive temporal correlation, whereas timepoints when one was active and the other inactive (or vice versa) would support a negative correlation. This can be quantified for all timepoints by first normalizing the timecourse of each region, subtracting the mean and dividing by the root sum-of-squares of the mean-centered data, and then taking the pointwise product of the two normalized timecourses (for more details, see Aly & Turk-Browne, 2016a). These products correspond directly to the contribution of each timepoint to the overall correlation — their sum *is* the Pearson correlation coefficient (Turk-Browne, 2013; Wang, Cohen, Li, & Turk-Browne, 2015; Worsley, Chen, Lerch, & Evans, 2005).

The result of this first step is a normalized product timecourse for each pair of ROIs in all attention runs (Figure 7A). To identify potential control regions, we then used this timecourse as a regressor in a GLM of voxelwise activity in the residuals of the ‘evoked’ model. We restricted analysis to frontal and parietal lobes (defined with MNI Structural Atlas) given the importance of frontoparietal cortex for top-down control (Corbetta and Shulman, 2002; Serences and Yantis, 2006; Noudoost et al., 2010) and to avoid circularity with the visual regions used to compute the product timecourses. The resulting parameter estimates indicate the extent to which each voxel’s activity was correlated with the products — that is, whether the region was more (or less) active during timepoints that increased or decreased background connectivity. A separate GLM was fit for each attention run (face-right, face-left, scene-right, scene-left) and pair of regions (left V4-FFA, right V4-FFA, left V4-PPA, and right V4-PPA). Of the 16 GLMs per subject, we sorted the resulting statistical maps by spatial and categorical relevance, mirroring the conditions in the main background connectivity analyses: space+/category+, space+/category-, space-/category+, space-/category-. These maps were combined at the group level within condition, treating subject as a random effect, with the reliability of each voxel’s correlation with the product timecourse compared against zero using the *randomise* function in FSL. We corrected for multiple comparisons using cluster-mass thresholding (cluster forming threshold *z* = 3.0).

## Results

### Behavior

Behavior in the repetition-detection task during the attention runs was generally fast and accurate (Figure 2A, 2B). Response times (RTs) for correctly detected repetitions were analyzed using a 2 (space: left vs. right) × 2 (category: faces vs. scenes) repeated-measures analysis of variance (ANOVA). This analysis revealed a significant main effect of space (F_(1, 14)_ = 5.55, *p* = 0.03), reflecting a left visual field advantage (Kimura, 1966). The main effect of category and the interaction were not significant (*p*s > 0.36).

**Figure 2.**
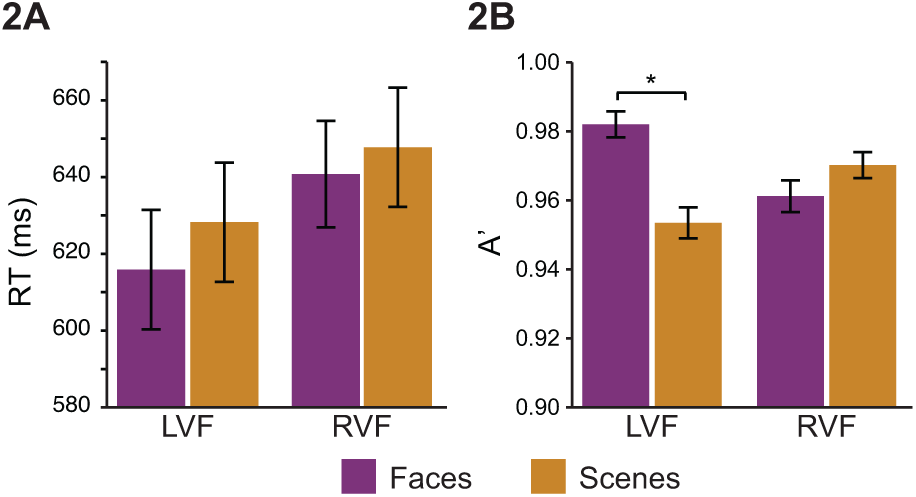
Behavioral results. (A) Participants were faster to respond to repetitions in the left visual field. (B) Participants were more accurate with face vs. scene repetitions in the LVF. *<.05. Error bars reflect +/- 1 SEM.

The accuracy of repetition detection was measured with *A’* (Grier, 1971). The same 2 (space) × 2 (category) ANOVA revealed a significant interaction (F_(1, 14)_ = 14.54, *p* = 0.002), but no reliable main effects of category (F_(1, 14)_ = 3.50, *p* = 0.08) or space (F_(1, 14)_ = 0.24, *p* = 0.63). The interaction reflects the fact that face repetitions were detected more accurately than scene repetitions in the left (*t*_(14)_ = 4.02, *p* = 0.001) but not right visual field (*t*_(14)_ = -1.24, *p* = 0.24), consistent with the particularly strong left visual field bias in face processing (De Renzi, Perani, Carlesimo, Silveri, & Fazio, 1994; Verosky & Turk-Browne, 2012; Yovel et al., 2008).

Because stimuli were presented 8° in the periphery, we were attuned to the possibility of systematic biases in eye position across conditions. We considered two measures — average horizontal displacement of gaze location from fixation and the percentage of samples more than 2° from fixation (i.e., the inner boundary of our stimuli) — and subjected each measure to a 2 (space) × 2 (category) ANOVA. Participants tended to look slightly to the right of fixation on average (*M* = +1.01°, *SD* = 0.78), but this bias did not differ across attention runs. Indeed, there were no reliable main effects or interactions for either measure (*p*s > 0.32).

### Evoked responses

Although our primary interest was attentional modulation of connectivity, we first examined the more conventional index of attention — the amplitude of the BOLD response in right and left V4, right FFA, and right PPA (Figure 3A). To quantify the amplitude of response (Figure 3B), we collapsed across ROIs and conditions and selected the time points across all blocks in which the BOLD percent signal change reliably differed from rest. This resulted in a set of ‘stimulated’ volumes (2–16; 1 and 17–24 were ‘non-stimulated’), over which we averaged the BOLD percent signal change within each ROI and condition.

**Figure 3.**
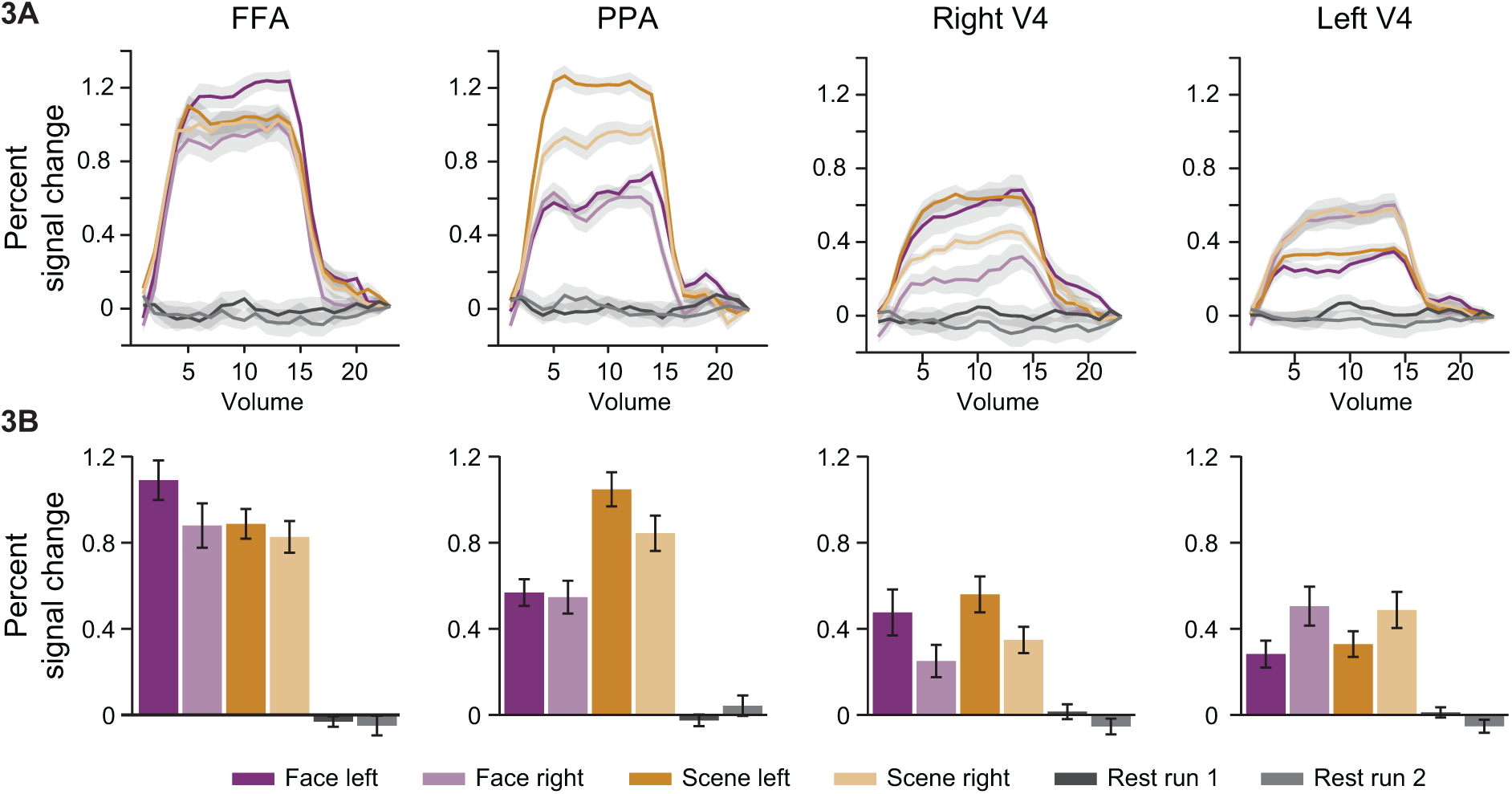
Evoked responses. (A) Timecourse of FIR parameter estimates for all conditions in every region, modeling the average evoked BOLD response. (B) Average FIR parameter estimates over ‘stimulated’ timepoints, as an index of response amplitude. Shaded regions and error bars reflect +/- 1 within-subject SEM.

First, we analyzed how V4 was modulated by attention. We computed a 2 (ROI: left/right V4) x 2 (spatial attention: left vs. right) x 2 (categorical attention: face vs. scene) ANOVA. We hypothesized that there would be a contralateral effect of spatial attention, which was supported by an interaction between ROI and spatial attention (F_(1,14)_ = 30.19, *P* < 0.001). This interaction was driven by greater BOLD response in right V4 for left vs. right attention runs (*t*_(14)_ = 3.51, *p* = 0.003) and in left V4 for right vs. left attention runs (*t*_(14)_ = 4.43, *p* < 0.001). This is consistent with the fact that V4 activity is modulated by spatial attention (Moran & Desimone, 1985). A complete report of the effects in this ANOVA can be found in Table 1A.

**Table 1.** **Evoked responses** (A) Modulation of right and left V4 by attention. (B). Modulation of FFA and PPA by attention. *indicates p < 0.05.

| Effect | DFn | DFd | F | p |
| --- | --- | --- | --- | --- |
| ROI | 1 | 14 | 0.02 | 0.90 |
| Space | 1 | 14 | 0.14 | 0.72 |
| Category | 1 | 14 | 2.15 | 0.16 |
| ROI x Space | 1 | 14 | 30.19 | *$<0.001$ |
| ROI x Category | 1 | 14 | 7.15 | *0.02 |
| Space x Category | 1 | 14 | 0.18 | 0.68 |
| ROI x Space x Category | 1 | 14 | 1.42 | 0.25 |

| Effect | DFn | DFd | F | p |
| --- | --- | --- | --- | --- |
| ROI | 1 | 14 | 6.74 | *0.02 |
| Space | 1 | 14 | 5.19 | *0.04 |
| Category | 1 | 14 | 9.92 | *0.007 |
| ROI x Space | 1 | 14 | 0.15 | 0.71 |
| ROI x Category | 1 | 14 | 45.31 | *$<0.001$ |
| Space x Category | 1 | 14 | 0.03 | 0.86 |
| ROI x Space x Category | 1 | 14 | 11.16 | *0.005 |

Next, we analyzed how ventral temporal regions were modulated by attention. We computed a 2 (ROI: FFA vs. PPA) x 2 (spatial attention: left vs. right) x 2 (categorical attention: face vs. scene) ANOVA. We hypothesized that the effect of categorical attention would depend on the selectivity of the regions, which was supported by an interaction between ROI and categorical attention (F_(1,14)_ = 45.31, *P* < 0.001). This interaction was driven by greater BOLD response in FFA for face vs. scene attention (*t*_(14)_ = 2.18, *p* = 0.05) and in PPA for scene vs. face attention (*t*_(14)_ = 7.19, *p <* 0.001). This is consistent with past findings that categorical attention modulates FFA and PPA (O’Craven et al., 1999).

This ANOVA also revealed that spatial attention modulated FFA and PPA. Specifically, there was a 3-way interaction between ROI, spatial attention, and categorical attention (F_(1,14)_ = 11.16, *p* = 0.005). This was driven by an increased BOLD response in FFA when attending to the left vs. right, but only for face attention (face attention: *t*_(14)_ = 2.21, *p* = 0.04; scene attention: *t*_(14)_ = 1.02, *p* = 0.33), and an increased BOLD response in PPA when attending to the left vs. right, but only for scene attention (face attention: *t*_(14)_ = 0.28, *p* = 0.78; scene attention: *t*_(14)_ = 2.41, *p* = 0.03). This might be explained by our ROI selection: The FFA could only be reliably defined in the right hemisphere, and we limited PPA to the right hemisphere to match (otherwise connectivity differences could reflect distance in the brain or hemispheric differences). Thus, the enhanced response for left attention (and preferred category) may reflect the slight contralateral bias of these ROIs (Hemond et al., 2007; MacEvoy & Epstein, 2007). A complete report of the effects in this ANOVA can be found in Table 1B.

### Background connectivity

We previously demonstrated that coupling between V4 and FFA/PPA is modulated by categorical attention (Al-Aidroos et al., 2012), and so first examined how categorical and spatial attention interacted to modulate coupling in the same regions. We investigated background connectivity between four pairs of ROIs: left V4 and FFA, right V4 and FFA, left V4 and PPA, and right V4 and PPA. We labeled these connections differently for each attention run, based on whether the V4 region coded for the attended location and whether the ventral temporal region coded for the attended category (Figure 4). This led to four connection types: both the V4 region and ventral temporal region coded for attended information (space+/category+), only the V4 region coded for attended information (space+/category-), only the ventral temporal region coded for attended information (space-/category+), and neither region coded for attended information (space-/category-). For example, when attending to faces on the right: left V4/FFA was labeled space+/category+, left V4/PPA was labeled space+/category-, right V4/FFA was labeled space-/category+, and right V4/PPA was labeled space-/category-.

**Figure 4.**
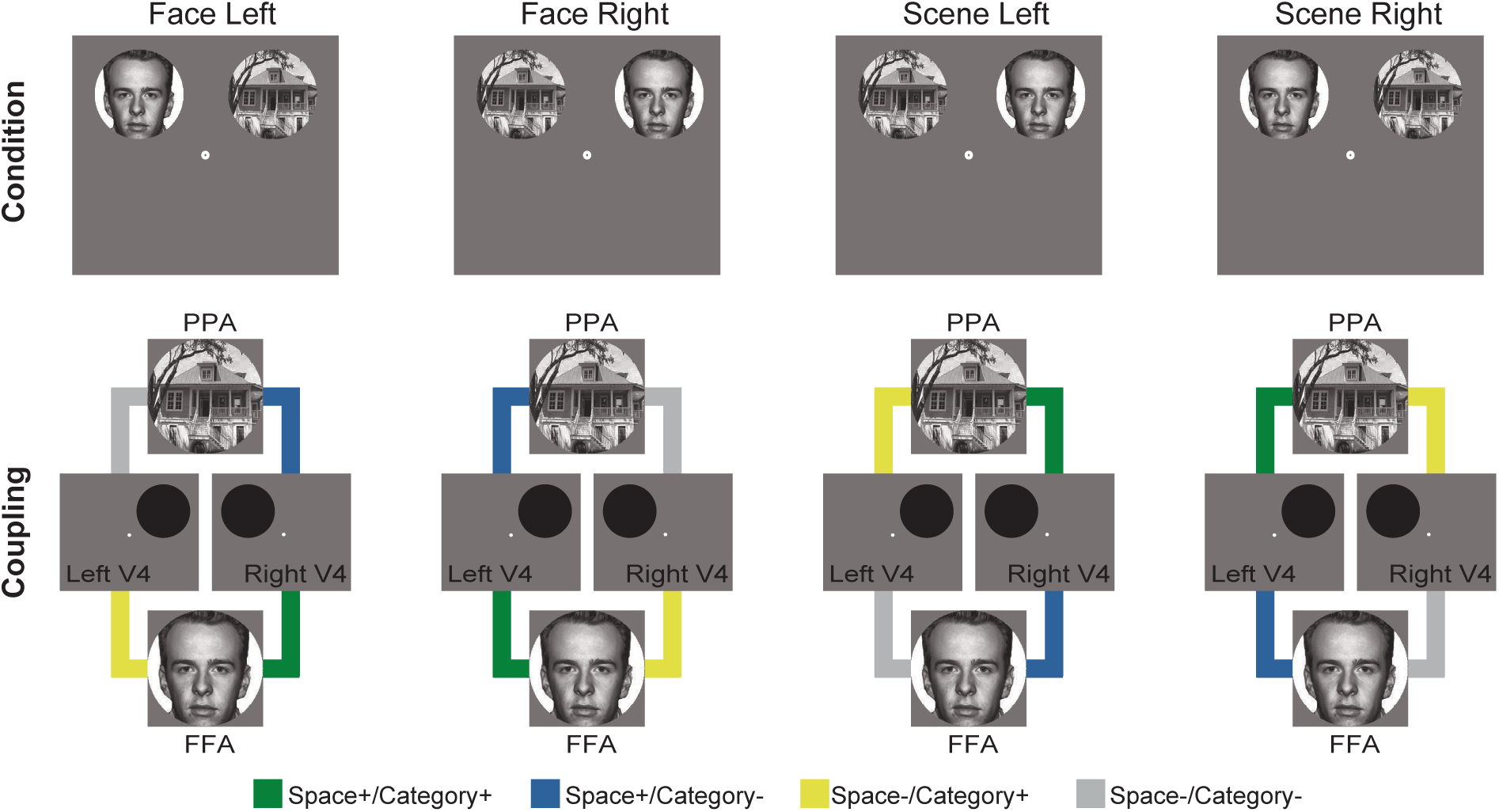
Connectivity types. Connectivity was measured between four pairs of ROIs: left V4 and FFA, right V4 and FFA, left V4 and PPA, and right V4 and PPA. These pairs were labeled based on the relevance of the constituent regions to the attended stimuli. Left and right V4 were relevant to right and left attention, respectively. FFA and PPA were relevant to face and scene attention, respectively.

This labeling resulted in a factorial design that allowed us to evaluate how categorical attention modulated the coupling of regions that code for spatially attended and unattended locations. Using a 2 (spatial relevance of connection: + vs. -) x 2 (categorical relevance of connection: + vs. -) ANOVA, we found a main effect of spatial relevance (*F*_(1, 14)_ = 17.25, *p* < 0.001), which was driven by greater connectivity when the V4 contralateral to the attended location was included in the region pair, both with the category-relevant ventral temporal ROI (space+/category+ > space-/category+: *t*_(14)_ = 5.21, *p* < 0.001) and marginally with the category-irrelevant ventral temporal ROI (space+/category-> space-/category-: *t*_(14)_ = 2.14, *p* = 0.05). In contrast, there was no main effect of categorical relevance (*F*_(1, 14)_ = 1.13, *p* = 0.31).

However, there was an effect of categorical relevance that depended on spatial relevance, as indicated by an interaction between these two factors (*F*_(1, 14)_ = 7.73, *p* = 0.01). This interaction was driven by greater background connectivity with the ventral temporal region that coded for the attended category relative to the unattended category, but only when paired with the V4 region that coded for the attended location (space+/category+ > space+/category-: *t*_(14)_ = 2.24, *p* = 0.04) and not the V4 region that coded for the unattended location (space-/category+ > space-/category+: *t*_(14)_ = -0.71, *p* = 0.49). Together these results suggest that the effects of categorical attention on background connectivity were spatially specific, enhancing connectivity only with the V4 that coded for the attended location.

### Intrinsic vs. evoked contributions to background connectivity

To what extent do these modulations reflect spontaneous internal interactions between cortical regions vs. correlations in BOLD activity evoked by external stimuli? The use of background connectivity above was an attempt to account for correlations driven by time-locked stimulus responses. However, it remains possible that our model did not perfectly capture all evoked responses. To assess whether these responses contributed to our results, we performed two further analyses.

We first evaluated whether background connectivity was modulated in the absence of evoked responses, during the ‘non-stimulated’ volumes (i.e., timepoints with responses that did not differ from baseline). The same 2 (spatial relevance of connection: + vs. -) x 2 (categorical relevance of connection: + vs. -) ANOVA restricted to these volumes revealed a main effect of spatial relevance (F_(1, 14)_ = 6.84, *p* = 0.02), no main effect of categorical relevance (F_(1, 14)_ = 0.87, *p* = 0.37), and no interaction (F_(1, 14)_ = 1.52, *p* =.24). From these results, we can conclude that the modulation of background connectivity by spatial attention reflects intrinsic activity. The lack of a reliable interaction leaves open the possibility that evoked activity contributed to the interaction reported above. However, this null effect should be interpreted with caution. For example, this analysis may have been underpowered because it only considered 9 volumes per block (as opposed to 24 in the main analysis). Therefore, we turned to another approach for assessing the contribution of evoked responses.

The same task structure was used for every participant, and thus evoked responses should be synchronized across participants (Hasson, Nir, Levy, Fuhrmann, & Malach, 2004). In contrast, intrinsic activity is idiosyncratic and we would not expect it to be synchronized. Thus, examining correlations across rather than within participants can help diagnose evoked vs. intrinsic activity. Specifically, if the observed modulation of background connectivity between V4 and FFA/PPA within participants reflects evoked activity, the same results should be obtained by correlating one participant’s V4 with other participants’ FFA/PPA. To perform this analysis, right and left V4 background timecourses from each participant were correlated with the average FFA and PPA background timecourses over all other participants, separately for every ROI pair and attention run. The 2 (spatial relevance of connection: + vs. -) x 2 (categorical relevance of connection: + vs. -) ANOVA applied to these across-participant correlations revealed no main effect of spatial relevance (F_(1, 14)_ = 0.42, *p* = 0.53), no main effect of categorical relevance (F_(1, 14)_ = 0.37, *p* = 0.55), and no interaction (F_(1, 14)_ = 1.88, *p* = 0.19). To verify that there were no alignment or other problems and that this analysis would have been sensitive to shared responses, the average correlation across conditions prior to removing the evoked activity in the background connectivity procedure was robust (mean *r* = 0.47; *t*_(14)_ = 11.82, *p* < 0.001), and reliably greater than in the resulting background data (mean *r* = 0.001; *t*_(14)_ = 0.15, *p* = 0.88; comparison: *t*_(14)_ = 11.43, *p* < 0.001). Thus, these timecourses contained neither across-participant correlations nor modulation of these correlations by attention. Critically, these same timecourses produced the main results when the correlations were calculated within participant (Fig. 5), emphasizing the role of intrinsic activity.

**Figure 5.**
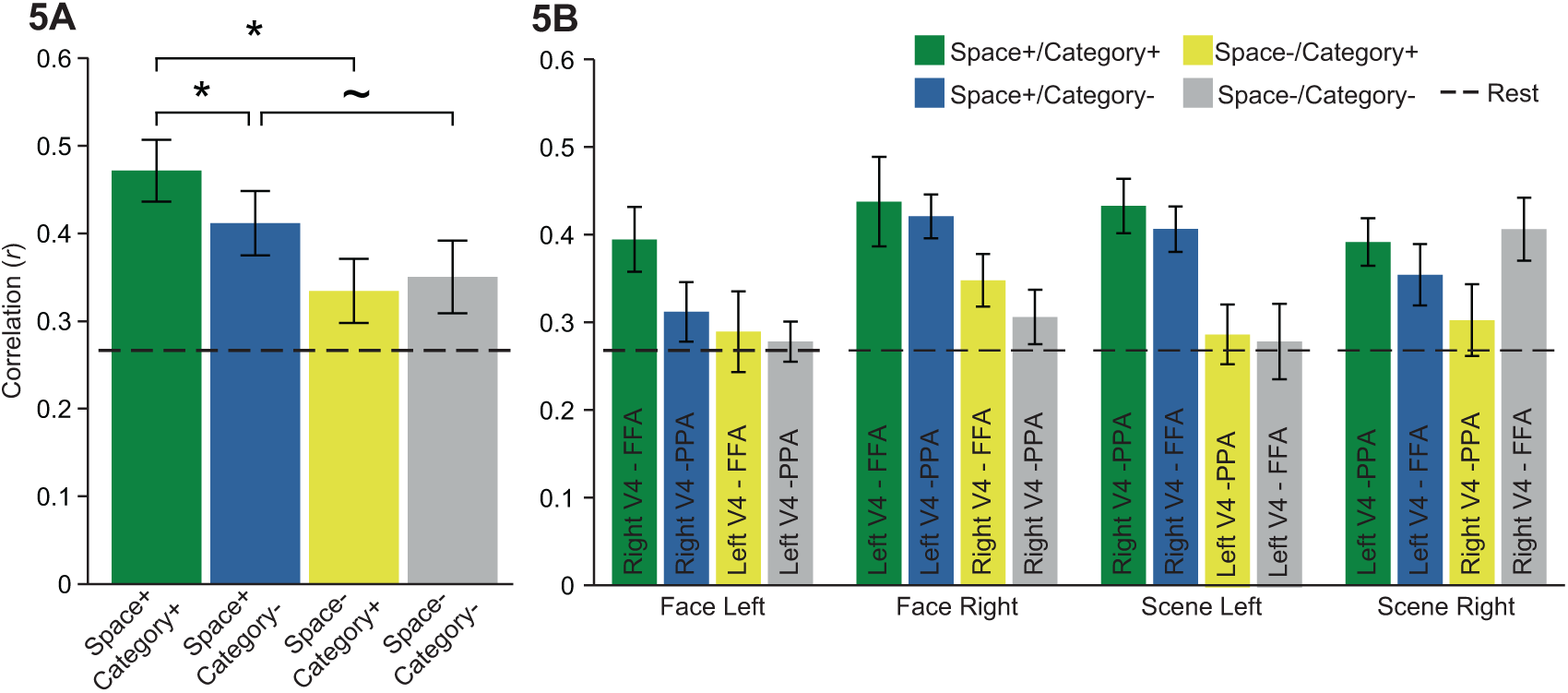
Background connectivity. (A) Spatial attention enhanced background connectivity between contralateral V4 and both FFA and PPA, while categorical attention modulated background connectivity of FFA and PPA only with the V4 region that coded for the attended spatial location. (B) Background connectivity in each attention run by spatial and categorical selectivity. *indicates *p* < 0.05, ∼ indicates *p* < 0.10. Error bars reflect within-subject SEM.

### Areas V1-V3

We next conducted exploratory analyses to assess whether modulation of connectivity by spatial attention is specific to V4, or if similar effects could be seen earlier in the visual stream. We repeated the main background connectivity analysis in retinotopic areas V1-V3. These areas can be found in both the ventral and dorsal streams, with selectivity for the upper and lower visual field, respectively. Because spatial attention was directed to the upper visual field (where images appeared), we focused on ventral areas (Figure 6A). We computed a 2 (spatial relevance of connection: + vs. -) x 2 (categorical relevance of connection: + vs. -) ANOVA for each ROI. We found no main effects of spatial or categorical relevance, nor any interactions in ventral V1, V2, or V3 (*Fs* < 2.31, *p*s > 0.15). Because the lower visual field was always spatially irrelevant, we reasoned that there might be different, possibly opposite effects in dorsal areas (Figure 6B). Indeed, there was a main effect of spatial relevance in dorsal V1 (*F* _(1, 14)_ = 7.22, *p* = 0.02) but not V2 or V3 (*Fs* < 0.26, *p* > 0.62), and main effects of categorical relevance in all three dorsal areas (V1: *F*_(1, 14)_ = 4.47, *p* = 0.05; V2: *F*_(1, 14)_ = 8.15, *p* = 0.01; V3: *F*_(1, 14)_ = 6.20, *p* = 0.03). Interestingly, and different from V4, these effects were driven by *lower* connectivity for relevant connections. There were no interactions in the dorsal ROIs (*Fs* < 2.89, *p* > 0.11).

**Figure 6.**
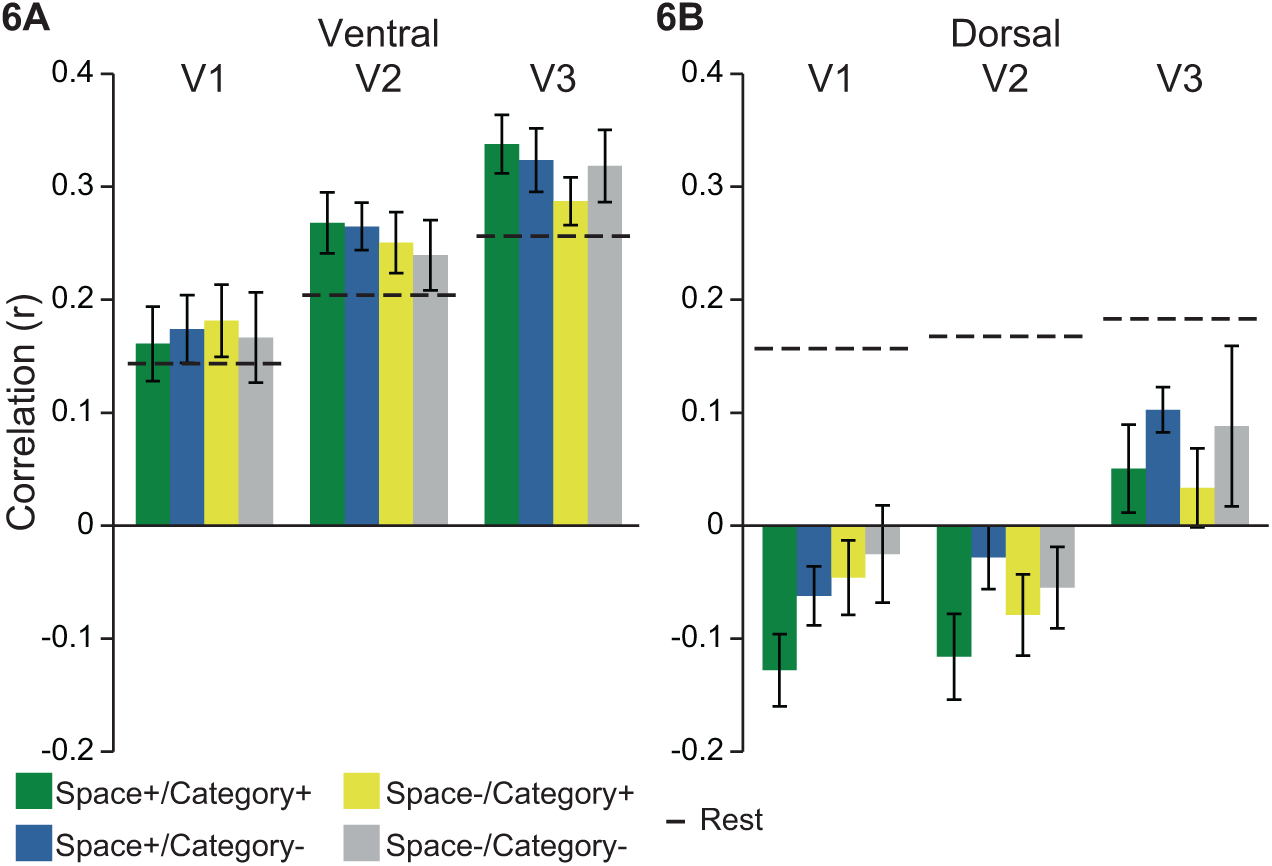
Background connectivity in V1, V2 and V3. (A) The interaction between spatial and categorical relevance in V4 did not extend into ventral V1-V3. (B) In the dorsal stream, background connectivity with V1, V2 and V3 was modulated by categorical relevance and V1 by spatial relevance, but in the opposite direction of V4, with *decreased* coupling to the category-selective ventral temporal region. Overall, attention to the upper visual field (all conditions) had diverging effects relative to rest, with relatively enhanced connectivity for ventral areas and suppressed connectivity for dorsal areas. Error bars reflect within-subject SEM.

To further explore how spatial attention to the upper and lower visual fields modulated connectivity, we compared overall background connectivity collapsed across the four conditions against the average of the two rest runs using a 3 (ROI: V1, V2, V3) × 2 (stream: ventral, dorsal) x 2 (state: attention vs. rest). There was a main effect of ROI (*F*_(2, 28)_ = 8.48, *p* = 0.001), with increasing background connectivity from V1 to V3, and a main effect of stream (*F*_(1, 14)_ = 38.41, *p* < 0.001), with stronger connectivity for ventral than dorsal streams; the main effect of state was not reliable (*F*_(1, 14)_ = 2.94, *p* = 0.11). Critically, there was a highly robust interaction between stream and state (*F*_(1, 14)_ = 57.49, *p* < 0.001), with background connectivity during attention higher than during rest in the ventral stream and lower in the dorsal stream. There was also an interaction between ROI and state (*F*_(2, 28)_ = 3.58, *p* = 0.04), with the difference between attention and rest runs decreasing from V1 to V3; no other interactions reached significance (*F*s > 2.62, *p*s > 0.08).

### Temporal MUD analysis

Our primary hypotheses concerned modulation of interactions within the visual system. However, this modulation likely results from the deployment of top-down control regulated elsewhere in the brain. Previous studies have demonstrated that frontoparietal cortex supports such control (Corbetta and Shulman, 2002; Serences and Yantis, 2006; Noudoost et al., 2010), including by communicating with visual areas (Chadick & Gazzaley, 2011; Gregoriou et al., 2009; Saalmann et al., 2007). To explore the role of frontoparietal cortex in the modulation of background connectivity within visual cortex, we searched for voxels in the frontal and parietal lobes whose activity at a given timepoint predicted how much that timepoint contributed to the background connectivity between visual areas (Figure 7A-B). By performing this analysis separately for each attention condition, we could evaluate which control structures support spatial attention, categorical attention, and their integration (Figure 7C; Table 2).

**Figure 7.**
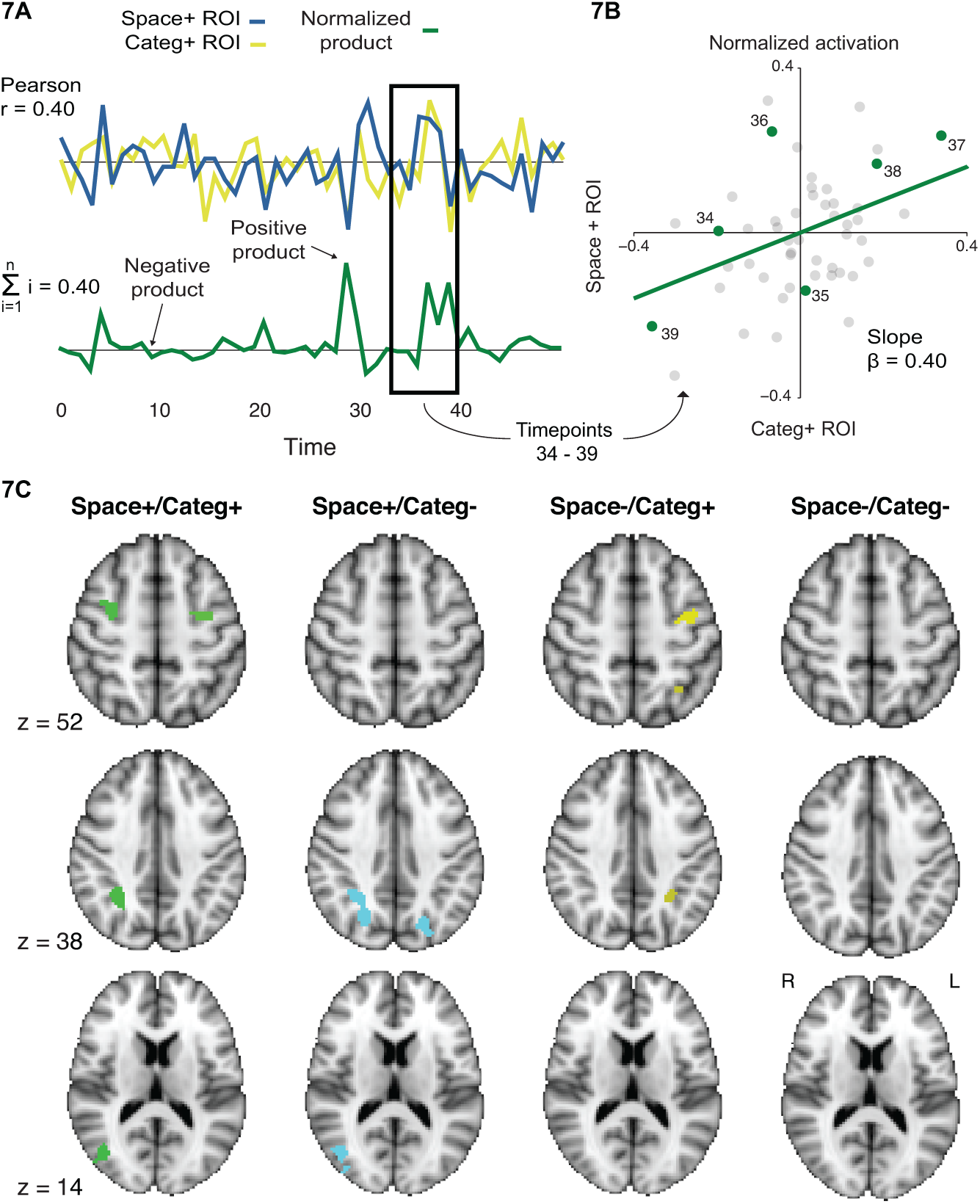
Temporal MUD analysis. (A) For each attention run, the normalized timecourses from right or left V4 were multiplied with the normalized timecourses from FFA or PPA. The resulting timecourse represented the contribution of each time point to the background connectivity between the two regions (the sum of the products is the Pearson correlation coefficient). (B) The graphical intuition for this analysis is that if both ROI timecourses have positive or negative normalized values at a given timepoint, then the timepoint falls in the 1^st^ or 3^rd^ quadrant of a scatterplot relating the normalized activity of one region to another. Because of the mean-centering of both regions, the best-fit line passes through origin and thus points in these quadrants support a positive slope. If one timecourse has a positive value and the other negative, then the timepoint falls in the 2^nd^ or 4^th^ quadrant, which supports a negative slope. The relative balance of points in quadrants 1/3 vs. 2/4 thus determines the sign of the slope, and because of the variance normalization, the value of the slope is the Pearson correlation coefficient. (C) When examining voxels whose BOLD timecourses correlated with the product timecourse for each condition, we observed significant clusters in the core attention network: bilateral FEF, bilateral IPS/SPL, and right TPJ. Contrasts corrected for multiple comparisons using cluster-mass thresholding (*p* < 0.05; cluster-forming threshold *z* = 3).

**Table 2.** **Temporal MUD results.** Frontoparietal regions whose activity correlates with background connectivity within visual cortex. R and L indicate right and left hemisphere. Extent is size of the clusters in voxels. X, Y, and Z coordinates indicate the center of gravity in MNI space (mm). *p* corresponds to corrected significance of cluster.

|  | Extent | X | Y | Z | *p* |
| --- | --- | --- | --- | --- | --- |
| Space+/Category+ |  |  |  |  |  |
| L FEF | 171 | -29.8 | -8.0 | 47.6 | 0.003 |
| R FEF | 122 | 33.4 | -2.0 | 50.1 | 0.010 |
| R IPS/SPL | 114 | 29.7 | -51.8 | 36.8 | 0.011 |
| R TPJ | 94 | 43.4 | -71.1 | 12.3 | 0.015 |
| Space+/Category- |  |  |  |  |  |
| R IPS/SPL | 339 | 25.2 | -61.4 | 40.0 | 0.001 |
| L IPS/SPL | 261 | -22.6 | -71.1 | 35.9 | 0.001 |
| R TPJ | 237 | 41.1 | -74.1 | 16.9 | 0.002 |
| Space-/Category+ |  |  |  |  |  |
| L FEF | 139 | -35.8 | -8.4 | 51.9 | 0.011 |
| L Anterior IPS/SPL | 108 | -28.5 | -51.5 | 40.7 | 0.020 |

In the space+/category+ condition — i.e., the spatially relevant V4 area paired with the categorically relevant ventral temporal area — there were four frontoparietal clusters whose moment-by-moment activity tracked variation in background connectivity (corrected *p* < 0.05): left and right superior frontal sulcus/precentral sulcus (frontal eye fields, FEF; Hutchison et al., 2012), right anterior intraparietal sulcus (IPS)/superior parietal lobule (SPL), and right temporal parietal junction (TPJ). These are the core areas of the dorsal and ventral attention networks (Corbetta & Shulman, 2002). In the space+/category-condition, there were three clusters: left and right IPS/SPL and right TPJ. In the space-/category+ condition, there were two clusters: left FEF and left anterior IPS/SPL. No clusters emerged in the space-/category-condition.

## Discussion

Based on prior work about feature-based attention, our key finding that categorical attention selectively modulated connectivity with spatially relevant regions may seem surprising. A number of studies have found that attending to a feature at one location enhances the response to that feature throughout the visual field (Chelazzi, Miller, Duncan, & Desimone, 1993; Chelazzi, Duncan, Miller, & Desimone, 1998; McAdams & Maunsell, 1999; Treue & Trujillo, 1999; Hayden & Gallant, 2009; Cohen & Maunsell, 2011; Andersen, Fuchs, & Müller, 2009; Bondarenko et al., 2012; Andersen et al., 2013). Furthermore, attention to motion or color in one location enhances the BOLD response to the same motion or color in a distant, unattended spatial location (Saenz et al., 2002). This spread of feature-based attention even extends to spatial locations with no stimulus present (Serences & Boynton, 2007). Behaviorally, top-down attention enhances sensitivity to features outside of the attended location as well (Liu & Hou, 2011; Liu & Mance, 2011; White & Carrasco, 2011). We might then have expected that attention to faces or scenes in the right (or left) hemifield would enhance FFA and PPA connectivity with both left and right V4, as opposed to the selectively contralateral effect we observed. One possible explanation for our results is that evoked and intrinsic signals fundamentally differ in their spatial specificity, with the former being spatially global and the latter being spatially local. Indeed, a prior fMRI study of feature-based attention found that intrinsic baseline shifts in anticipation of a stimulus were restricted to the expected location of the stimulus, whereas V4 responses evoked by the stimulus were modulated at both expected and unexpected locations (McMains et al., 2007). In our case, rather than focusing on a pre-stimulus period, we isolated intrinsic signals by regressing out stimulus-evoked responses prior to calculating background connectivity.

Competition is another factor that may explain this finding (Moran & Desimone, 1985). Several studies have found enhanced responses to attended features in unattended locations only when there are competing stimuli within or outside of the attended spatial location (Saenz, Buraĉas, & Boynton, 2003; Zhang & Luck, 2009; Painter, Dux, Travis, & Mattingley, 2014), although others have found such effects regardless of competition (Bartsch et al., 2015). In our experiment, the stimuli in the unattended location were a different category than those in the attended location (e.g., faces on the left while attending to scenes on the right), and likely did not compete with the attended category/location. The lack of competing category may minimize any global enhancements in BOLD signal or correlated activity that would be expected otherwise. Furthermore, behaviorally, spatial attention and feature-based attention seem to operate independently except when unattended distractors are competitive with attended targets (Leonard, Balestreri, & Luck, 2015; White, Rolfs, & Carrasco, 2015). This suggests that while feature-based attention may globally enhance neural responses across the visual field, there may be later cognitive and neural processes that integrate over spatial and feature-based attention when needed. Switches in coupling between different levels of the visual system may be one such process.

This finding is also consistent with our prior study that found effects of categorical attention on background connectivity but did not consider spatial attention (Al-Aidroos et al., 2012). In that study, face and scene stimuli were overlaid in the same spatial location and fixated centrally. Bilateral V4 showed stronger background connectivity with FFA when faces were attended and with PPA when scenes were attended. Because both right and left V4 code for foveal stimuli, both regions were spatially relevant for the task. Thus, V4-FFA when faces were attended and V4-PPA when scenes were attended correspond to space+/category+ in the current study, and V4-PPA when faces were attended and V4-FFA when scenes were attended correspond to space+/category- in the current study. The fact that we found stronger connectivity for space+/category+ vs. space+/category- in the current study thus replicates these prior findings. The current study also extends these findings, by testing for effects of categorical attention on background connectivity with regions that do not code for the attended location (space-/category+ and space-/category-). That such effects were eliminated in unattended locations provides new insight into how attention modulates connectivity in the visual system, and into how different varieties of attention are integrated in the brain.

In this experiment, we operationalized coupling between regions as correlations in timecourses between two regions. However, correlated activity does not necessarily imply direct coupling or communication, and instead could reflect other extrinsic or stimulus-driven sources of correlations. We attempted to minimize some of these other sources by regressing out signals related to stimulus timing, head motion, and nuisance regions before calculating the correlations. Past work has found that this procedure gives rise to correlated activity – or ‘background connectivity’ – that is not driven by variance in BOLD signal across blocks or runs, another source of stimulus-driven correlations (Al-Aidroos et al., 2012). In that work, as well as in the present experiment, differences in intrinsic, subject-specific signals, rather than stimulus timing, underpin the observed modulation in background connectivity by attention. Such intrinsic signals still may not signify direct coupling between two regions, but instead reflect coordinated activity moderated by a third region. However, this does not necessarily undercut our findings – rather, it raises interesting questions about the broader network of control regions modulating correlations within visual cortex. Indeed, models of cognitive control emphasize the role of control regions in establishing sensorimotor pathways (Miller & Cohen, 2001). We explore these questions with an exploratory analysis that identified regions whose BOLD signal tracked correlations between two visual regions (see below for further discussion).

One potential drawback to our design is that, although participants were instructed to attend a specific category and spatial location, participants could potentially complete the task by using only category or spatial attention. An ideal design would use composite images of faces and houses in both locations to necessitate both forms of attention. However, through behavioral piloting we found that such a task was too difficult. Furthermore, any idiosyncratic strategies that participants might use in our current design would only work against finding interactions between categorical attention and spatial attention. In other words, we found that background connectivity was modulated by both spatial and categorical attention despite the fact that only one form of attention as necessary to perform the task.

Interestingly, we also found a broad decrease in background connectivity for dorsal visual areas during attention runs. One interpretation of this finding is that connectivity was suppressed because the lower visual field, for which these areas code, was task irrelevant. Future studies that compare attention to upper vs. lower visual fields would be needed to definitively support this conclusion. Nevertheless, as an additional piece of evidence, similar suppression of dorsal connectivity with FFA/PPA was not observed in our prior study in which the stimuli were foveated — in fact, and opposite from the current study, connectivity was enhanced relative to rest (Al-Aidroos et al., 2012, Fig. 2C).

Although we primarily focused on how coupling within visual cortex supports combined spatial and non-spatial attention, we also conducted an exploratory analysis of control mechanisms that may drive such coupling. Prior studies have observed that frontoparietal areas couple with visual areas when they are relevant for current attentional goals (Baldauf & Desimone, 2014; Chadick & Gazzaley, 2011; Griffis et al., 2015). For example, we previously found frontoparietal regions that correlated with FFA when faces were attended and with PPA when scenes were attended (Al-Aidroos et al., 2012). Here we adopted a different approach, seeking to identify frontoparietal areas that track variance in the *relationship* (i.e., connectivity) between visual areas rather than in the activity of individual areas. This may provide additional leverage for understanding how top-down control signals modulate network activity within visual cortex, and especially how different components of attention are integrated. For example, right FEF was active during moments of high connectivity, but only between areas that were spatially and categorically relevant (e.g., left V4 and FFA, respectively, when attending to faces on the right). In contrast, right IPS/SPL and TPJ also tracked connectivity in the visual system but only required spatial attention (e.g., left V4 and PPA in the example above), and left FEF tracked connectivity but only required categorical attention (e.g., right V4 and FFA). These findings lead to the prediction for future studies that right FEF in particular may play an important role in integrating attentional goals.

Beyond attentional modulation of background connectivity between V4 and FFA/PPA, we also observed stimulus-evoked responses in these regions. Specifically, spatial attention but not categorical attention modulated responses in right and left V4, whereas both spatial and categorical attention modulated responses in FFA and PPA. In examining connectivity, we took care to control for these evoked responses by removing them from the data. The fact that attentional modulation of background connectivity persisted into fixation periods without stimuli and did not synchronize across participants (despite identical stimulus timing) further suggests that our connectivity results do not depend on evoked responses. Nevertheless, these two mechanisms must interact to support top-down attention. One possibility is that good communication between regions may strengthen evoked responses downstream (Fries, 2005). Relatedly, connectivity may filter which information gets transmitted and lead to more selective processing in broadly tuned regions (Córdova, Tompary, & Turk-Browne, 2016). Here, connectivity with spatially selective V4 may aid FFA and PPA in processing category information from specific locations. Regardless, by focusing on covariance in neural activity typically ignored in analyses of evoked responses, our findings illustrate the distributed and integrative nature of attention.

## Acknowledgements

This work was supported by NSF GRFP (AT), NSERC PDF and DG (NA), and NIH R01 EY021755 (NTB).

## Notes

Conflict of interest: The authors declare no competing financial interests

## References

Al-Aidroos, N., Said, C. P., & Turk-Browne, N. B. (2012). Top-down attention switches coupling between low-level and high-level areas of human visual cortex. Proceedings of the National Academy of Sciences of the United States of America, 109(36), 14675–14680.

Aly, M., & Turk-Browne, N. B. (2016a). Attention promotes episodic encoding by stabilizing hippocampal representations. Proceedings of the National Academy of Sciences, 113(4), E420–E429.

Aly, M., & Turk-Browne, N. B. (2016b). Attention stabilizes representations in the human hippocampus. Cerebral Cortex, 26, 783–796.

Andersen, S. K., Fuchs, S., & Müller, M. M. (2009). Effects of feature-selective and spatial attention at different stages of visual processing. Journal of Cognitive Neuroscience, 23(1), 238–246.

Andersen, S. K., Hillyard, S. A., & Müller, M. M. (2013). Global facilitation of attended features is obligatory and restricts divided attention. Journal of Neuroscience, 33(46), 18200–18207.

Arcaro, M. J., McMains, S. A., Singer, B. D., & Kastner, S. (2009). Retinotopic organization of human ventral visual cortex. The Journal of Neuroscience, 29(34), 10638–10652.

Baldauf, D., & Desimone, R. (2014). Neural mechanisms of object-based attention. Science (New York, N.Y.), 344(6182), 424–427.

Bartsch, M. V., Boehler, C. N., Stoppel, C. M., Merkel, C., Heinze, H.-J., Schoenfeld, M. A., & Hopf, J.-M.(2015). Determinants of global color-based selection in human visual cortex. Cerebral Cortex, 25(9), 2828–2841.

Bondarenko, R., Boehler, C. N., Stoppel, C. M., Heinze, H.-J., Schoenfeld, M. A., & Hopf, J.-M. (2012). Separable mechanisms underlying global feature-based attention. Journal of Neuroscience, 32(44), 15284–15295.

Bosman, C. A., Schoffelen, J.-M., Brunet, N., Oostenveld, R., Bastos, A. M., Womelsdorf, T. Fries, P. (2012). Attentional stimulus selection through selective synchronization between monkey visual areas. Neuron, 75(5), 875–888.

Chadick, J. Z., & Gazzaley, A. (2011). Differential coupling of visual cortex with default or frontal-parietal network based on goals. Nature Neuroscience, 14(7), 830–832.

Chelazzi, L., Duncan, J., Miller, E. K., & Desimone, R. (1998). Responses of neurons in inferior temporal cortex during memory-guided visual search. Journal of Neurophysiology, 80(6), 2918–2940.

Chelazzi, L., Miller, E. K., Duncan, J., & Desimone, R. (1993). A neural basis for visual search in inferior temporal cortex. Nature, 363(6427), 345–347.

Cohen, E. H., & Tong, F. (2015). Neural mechanisms of object-based attention. Cerebral Cortex, 25(4), 1080–1092.

Cohen, M. R., & Maunsell, J. H. R. (2011). Using neuronal populations to study the mechanisms underlying spatial and feature attention. Neuron, 70(6), 1192–1204.

Connor, C. E., Gallant, J. L., Preddie, D. C., & Essen, D. C. V. (1996). Responses in area V4 depend on the spatial relationship between stimulus and attention. Journal of Neurophysiology, 75(3), 1306– 1308.

Corbetta, M., Miezin, F. M., Dobmeyer, S., Shulman, G. L., & Petersen, S. E. (1990). Attentional modulation of neural processing of shape, color, and velocity in humans. Science, 248(4962), 1556.

Corbetta, M., & Shulman, G. L. (2002). Control of goal-directed and stimulus-driven attention in the brain. Nature Reviews Neuroscience, 3(3), 201–215.

Córdova, N. I., Tompary, A., & Turk-Browne, N. B. (2016). Attentional modulation of background connectivity between ventral visual cortex and the medial temporal lobe. Neurobiology of Learning and Memory, 134, Part A, 115–122.

Dale, A. M., Fischl, B., & Sereno, M. I. (1999). Cortical surface-based analysis: I. Segmentation and surface reconstruction. NeuroImage, 9(2), 179–194.

De Renzi, E., Perani, D., Carlesimo, G. A., Silveri, M. C., & Fazio, F. (1994). Prosopagnosia can be associated with damage confined to the right hemisphere—An MRI and PET study and a review of the literature. Neuropsychologia, 32(8), 893–902.

Duncan, K., Tompary, A., & Davachi, L. (2014). Associative encoding and retrieval are predicted by functional connectivity in distinct hippocampal area CA1 pathways. Journal of Neuroscience, 34(34), 11188–11198.

Fischl, B., Sereno, M. I., & Dale, A. M. (1999). Cortical surface-based analysis: II: Inflation, flattening, and a surface-based coordinate system. NeuroImage, 9(2), 195–207.

Fox, M. D., & Raichle, M. E. (2007). Spontaneous fluctuations in brain activity observed with functional magnetic resonance imaging. Nature Reviews Neuroscience, 8(9), 700–711.

Fries, P. (2005). A mechanism for cognitive dynamics: neuronal communication through neuronal coherence. Trends in Cognitive Sciences, 9(10), 474–480.

Furey, M. L., Tanskanen, T., Beauchamp, M. S., Avikainen, S., Uutela, K., Hari, R., & Haxby, J. V. (2006). Dissociation of face-selective cortical responses by attention. Proceedings of the National Academy of Sciences of the United States of America, 103(4), 1065–1070.

Giesbrecht, B., Woldorff, M. G., Song, A. W., & Mangun, G. R. (2003). Neural mechanisms of top-down control during spatial and feature attention. NeuroImage, 19(3), 496–512.

Goldsmith, M., & Yeari, M. (2003). Modulation of object-based attention by spatial focus under endogenous and exogenous orienting. Journal of Experimental Psychology: Human Perception and Performance, 29(5), 897–918.

Gregoriou, G. G., Gotts, S. J., Zhou, H., & Desimone, R. (2009). High-frequency, long-range coupling between prefrontal and visual cortex during attention. Science, 324(5931), 1207–1210.

Grier, J. B. (1971). Nonparametric indexes for sensitivity and bias: computing formulas. Psychological Bulletin, 75(6), 424.

Griffis, J. C., Elkhetali, A. S., Burge, W. K., Chen, R. H., & Visscher, K. M. (2015). Retinotopic patterns of background connectivity between V1 and fronto-parietal cortex are modulated by task demands. Frontiers in Human Neuroscience, 338.

Hasson, U., Nir, Y., Levy, I., Fuhrmann, G., & Malach, R. (2004). Intersubject synchronization of cortical activity during natural vision. Science, 303, 1634–1640.

Hayden, B. Y., & Gallant, J. L. (2009). Combined effects of spatial and feature-based attention on responses of V4 neurons. Vision Research, 49(10), 1182–1187.

Hemond, C. C., Kanwisher, N. G., & Op de Beeck, H. P. (2007). A preference for contralateral stimuli in human object- and face-selective cortex. PLoS ONE, 2(6), e574.

Hutchison, R. M., Gallivan, J. P., Culham, J. C., Gati, J. S., Menon, R. S., & Everling, S. (2012). Functional connectivity of the frontal eye fields in humans and macaque monkeys investigated with resting-state fMRI. Journal of Neurophysiology, 107(9), 2463–2474.

Kimura, D. (1966). Dual functional asymmetry of the brain in visual perception. Neuropsychologia, 4.

Leonard, C. J., Balestreri, A., & Luck, S. J. (2015). Interactions between space-based and feature-based attention. Journal of Experimental Psychology: Human Perception and Performance, 41(1), 11–16.

Liu, T., & Hou, Y. (2011). Global feature-based attention to orientation. Journal of Vision, 11(10), 8–8.

Liu, T., & Mance, I. (2011). Constant spread of feature-based attention across the visual field. Vision Research, 51(1), 26–33.

MacEvoy, S. P., & Epstein, R. A. (2007). Position selectivity in scene- and object-responsive occipitotemporal regions. Journal of Neurophysiology, 98(4), 2089–2098.

McAdams, C. J., & Maunsell, J. H. R. (1999). Effects of attention on orientation-tuning functions of single neurons in macaque cortical area V4. The Journal of Neuroscience, 19(1), 431–441.

McMains, S. A., Fehd, H. M., Emmanouil, T.-A., & Kastner, S. (2007). Mechanisms of feature- and space- based attention: Response modulation and baseline increases. Journal of Neurophysiology, 98(4), 2110–2121.

Miller, E. K., & Cohen, J. D. (2001). An integrative theory of prefrontal cortex function. Annual Review of Neuroscience, 24(1), 167–202.

Moran, J., & Desimone, R. (1985). Selective attention gates visual processing in the extrastriate cortex. Science (New York, N.Y.), 229(4715), 782–784.

Motter, B. C. (1993). Focal attention produces spatially selective processing in visual cortical areas V1, V2, and V4 in the presence of competing stimuli. Journal of Neurophysiology, 70(3), 909–919.

Müller, N. G., & Kleinschmidt, A. (2003). Dynamic interaction of object- and space-based attention in retinotopic visual areas. The Journal of Neuroscience, 23(30), 9812–9816.

Norman-Haignere, S. V, McCarthy, G., Chun, M. M., & Turk-Browne, N. B. (2012). Category-selective background connectivity in ventral visual cortex. Cerebral Cortex, 22(2), 391–402.

Nyström, M., & Holmqvist, K. (2010). An adaptive algorithm for fixation, saccade, and glissade detection in eyetracking data. Behavior Research Methods, 42(1), 188–204.

O’Craven, K. M., Downing, P. E., & Kanwisher, N. (1999). fMRI evidence for objects as the units of attentional selection. Nature, 401(6753), 584–587.

Painter, D. R., Dux, P. E., Travis, S. L., & Mattingley, J. B. (2014). Neural responses to target features outside a search array are enhanced during conjunction but not unique-feature search. The Journal of Neuroscience, 34(9), 3390–3401.

Saalmann, Y. B., Pigarev, I. N., & Vidyasagar, T. R. (2007). Neural mechanisms of visual attention: How top-down feedback highlights relevant locations. Science, 316(5831), 1612–1615.

Saenz, M., Buracas, G. T., & Boynton, G. M. (2002). Global effects of feature-based attention in human visual cortex. Nature Neuroscience, 5(7), 631–632.

Saenz, M., Buraĉas, G. T., & Boynton, G. M. (2003). Global feature-based attention for motion and color. Vision Research, 43(6), 629–637.

Schwarzlose, R. F., Swisher, J. D., Dang, S., & Kanwisher, N. (2008). The distribution of category and location information across object-selective regions in human visual cortex. Proceedings of the National Academy of Sciences of the United States of America, 105(11), 4447–4452.

Serences, J. T., & Boynton, G. M. (2007). Feature-based attentional modulations in the absence of direct visual stimulation. Neuron, 55(2), 301–312.

Serences, J. T., Schwarzbach, J., Courtney, S. M., Golay, X., & Yantis, S. (2004). Control of object-based attention in human cortex. Cerebral Cortex, 14(12), 1346–1357.

Shomstein, S., & Yantis, S. (2004). Control of attention shifts between vision and audition in human cortex. The Journal of Neuroscience, 24(47), 10702–10706.

Shomstein, S., & Yantis, S. (2006). Parietal cortex mediates voluntary control of spatial and nonspatial auditory attention. The Journal of Neuroscience, 26(2), 435–439.

Smith, S. M., Jenkinson, M., Woolrich, M. W., Beckmann, C. F., Behrens, T. E.., Johansen-Berg, H., others. (2004). Advances in functional and structural MR image analysis and implementation as FSL. NeuroImage, 23, S208–S219.

Soto, D., & Blanco, M. J. (2004). Spatial attention and object-based attention: A comparison within a single task. Vision Research, 44(1), 69–81.

Summerfield, C., Greene, M., Wager, T., Egner, T., Hirsch, J., & Mangels, J. (2006). Neocortical connectivity during episodic memory formation. PLoS Biology, 4(5), e128.

Tompary, A., Duncan, K., & Davachi, L. (2015). Consolidation of associative and item memory is related to post-encoding functional connectivity between the ventral tegmental area and different medial temporal lobe subregions during an unrelated task. The Journal of Neuroscience, 35(19), 7326–7331.

Treue, S., & Trujillo, J. C. M. (1999). Feature-based attention influences motion processing gain in macaque visual cortex. Nature, 399(6736), 575–579.

Turk-Browne, N. B. (2013). Functional interactions as big data in the human brain. Science, 342(6158), 580–584.

Verosky, S. C., & Turk-Browne, N. B. (2012). Representations of facial identity in the left hemisphere require right hemisphere processing. Journal of Cognitive Neuroscience, 24(4), 1006–1017.

Wandell, B. A., Dumoulin, S. O., & Brewer, A. A. (2007). Visual field maps in human cortex. Neuron, 56(2), 366–383.

Wang, Y., Cohen, J. D., Li, K., & Turk-Browne, N. B. (2015). Full correlation matrix analysis (FCMA): An unbiased method for task-related functional connectivity. Journal of Neuroscience Methods, 251, 108–119.

White, A. L., & Carrasco, M. (2011). Feature-based attention involuntarily and simultaneously improves visual performance across locations. Journal of Vision, 11(6), 15–15.

White, A. L., Rolfs, M., & Carrasco, M. (2015). Stimulus competition mediates the joint effects of spatial and feature-based attention. Journal of Vision, 15 (14), 7–7.

Worsley, K. J., Chen, J.-I., Lerch, J., & Evans, A. C. (2005). Comparing functional connectivity via thresholding correlations and singular value decomposition. Philosophical Transactions of the Royal Society of London B: Biological Sciences, 360(1457), 913–920.

Yovel, G., Tambini, A., & Brandman, T. (2008). The asymmetry of the fusiform face area is a stable individual characteristic that underlies the left-visual-field superiority for faces. Neuropsychologia, 46(13), 3061–3068.

Zhang, W., & Luck, S. J. (2009). Feature-based attention modulates feedforward visual processing. Nature Neuroscience, 12(1), 24–25.

Zhou, H., & Desimone, R. (2011). Feature-based attention in the frontal eye field and area V4 during visual search. Neuron, 70(6), 1205–1217.

